# Constraints on the deformation of the vibrissa within the follicle

**DOI:** 10.1101/2020.04.20.050757

**Authors:** Yifu F. Luo, Chris S. Bresee, John W. Rudnicki, Mitra J. Z. Hartmann

## Abstract

Nearly all mammals have a vibrissal system specialized for tactile sensation, composed of whiskers growing from sensor-rich follicles in the skin. Because a whisker has no sensors along its length, an open question is how mechanoreceptors in the follicle transduce sensory signals. These mechanoreceptors are activated by whisker deflection, so it is essential to understand how the whisker deforms within the follicle and so how it may activate different populations of mechanoreceptors in different ways. During active whisking behaviors, muscle contractions and increases in blood pressure in the ring sinus will likely affect the whisker deformation profile. Directly recording from mechanoreceptors under these conditions is difficult due to their small size, location within intricate and delicate membranes, and movement during sensation. Using data from a previous experimental study on whisker deflection, and from histological analysis of follicle tissue, we develop a mechanical model of the follicle sinus complex. With this model we first simulate passive whisker contact, replicating previous results of *ex vivo* experiments on deformation of a whisker within the follicle. We then simulate whisker deformation within the follicle during active whisking. Results of these simulations predict that both intrinsic muscle contraction and elevated hydrostatic pressure within the ring sinus may be regulatory mechanisms to enhance tactile sensitivity during active whisking. The mechanical model presented in this study is an important first step in simulating mechanical interactions within whisker follicles, and aids in the development of artificial robotic follicles.

**Author summary:** Many mammals rely on whiskers as a mode of tactile sensation, especially when exploring in darkness. Active, rhythmic protraction and retraction of the whiskers, commonly referred to as whisking, is observed among many whisker specialist animals. Whisker-based sensing requires the forces and moments generated by external stimuli to be transduced into neural signals inside the follicle, which holds the base of the whisker shaft. Within the follicle, the interaction between the whisker’s deformation and the surrounding tissue determines how different groups of mechanoreceptors along the inside of a follicle will deformed. However, experimental measurement of this interaction is challenging to perform in active animals. We therefore created a mechanical model for the follicle sinus complex to simulate whisker deformation within the follicle resulting from external whisker deflection. Our simulations replicate results from previous *ex vivo* experiments that have monitored whisker deformation in the follicle ring sinus. We extend these results by predicting whisker deformation profiles during active whisking. Our results suggest that both intrinsic muscle contraction and an increase in blood pressure will affect the whisker deformation profile within the follicle, and in turn, the tactile sensitivity of the whisker system.

## Introduction

Nearly all mammals have a vibrissal (whisker) system [1], with some being specialized to actively gather tactile information from the environment [2–4]. Unlike an insect antenna, a whisker has no mechanoreceptors along its length. Instead, external mechanical stimuli are transmitted to a richly innervated follicle at the whisker base [5–9], where mechanoreceptors transduce the mechanical information into electrical signals [10, 11].

Follicle anatomy has been found to share a strikingly similar structure for multiple species across a broad swath of the mammalian family tree. Among these species are the rakali (*Hydromys chrysogaster*) [12], naked mole-rats (*Heterocephalus glaber*) [13], common mole rats (*Cryptomys hottentotus*) [14], tree squirrels (*Sciurus vulgaris*) [15], shrews (*Sorex araneus*) [16], rock hyrax (*Procavia capensis*) [17], tammar wallaby (*Macropus eugenii*) [18], manatee (*Trichechus manatus*) [19, 20], harbor seal (*Phoca vitulina*) [21], ringed seal (*Pusa hispida*) [22], California sea lion (*Zalophus californianus*) [23], sea otter (*Enhydra lutris*) [24], bearded seal (*Erignathus barbatus*) [25], Eurasian otter and pole cat (*Lutra lutra* and *Mustela putorius*) [26], rats and cats (*Rattus norvegicus* and *Felis catus*) [10]. When sectioned lengthwise, the cross-section of each follicle tends to be near-cylindrical (bearded-seal and shrew), ovular (rat and cat), or might resemble an inverted vase (squirrel). Regardless of shape, all follicles are densely packed with mechanoreceptors, often including Merkel, lanceolate, and club-like endings, and all contain one or more blood sinuses, which have been postulated to help regulate sensor sensitivity based on variations in blood pressure [27, 28].

An animal’s perception of a tactile stimulus will be determined by how these mechanoreceptors transduce mechanical deformation into neural signals, which will in turn be determined by how the vibrissa deforms within the follicle. Additionally, the particular profile of whisker deformation has the potential to actuate different populations of mechanoreceptors along the length of the follicle. We therefore investigate the deformation profile of the whisker within the follicle.

To begin to quantify the deformation of the whisker within the follicle, Whiteley et al. [29] recently performed a set of experiments to determine how the internal follicle tissue at the ring sinus (RS) level deformed in response to a vibrissal deflection. These experiments provide ground truth for tissue displacement resulting from passive whisker deformation relative to the dissected follicle, and so provide a starting point for predicting more complex whisker-follicle interactions.

In the present study, we develop a mechanical model of the whisker follicle sinus complex (FSC) that replicates the deformation profile observed in passive conditions [29]. We then use the model to predict the deformation profile during active whisking and examine the possible effects of blood pressure variation within the sinus. Our results suggest that active muscle contraction, as well as blood pressure, increases during the arousal concomitant to active exploration may both help enhance tactile sensitivity.

## Materials and Methods

Ethics Statement: All experiments involving animals were approved in advance by the Institutional Animal Care and Use Committee of Northwestern University.

### Anatomical experiments to estimate tissue stiffness along the follicle length

To obtain estimates of tissue stiffness within the follicle, we sectioned four mystacial pads of three adult (3 – 8 months), female, Long Evans rats (*Rattus norvegicus*). After use in unrelated electrophysiology experiments, rats were perfused with 1x phosphate-buffered saline solution (PBS) with 10 units/ml heparin and then with HistoChoice™. The mystacial pad tissue was dissected away from the underlying bone and placed in 100% HistoChoice™ overnight. After 24 hours, tissue was sequentially cryoprotected in 10%, 20%, and 30% sucrose in PBS, each until osmotic pressure was equalized, as indicated by the tissue resting on the bottom of the vial. Tissue was then flash-frozen in Optimal Cutting Temperature compound (Tissue-Tek^®^ O.C.T., Sakura Finetek) on a level aluminum block partially submerged in liquid nitrogen, and sectioned at 20 microns on an upright freezing microtome.

Tissue sections were mounted on gelatin coated slides using a 4% paraformaldehyde solution for 15 min, and then permeabilized with acetone for 5 min. Sections were washed, bleached, and stained in Mallory's Phosphotungstic Acid Hematoxilin (PTAH), washed again and dehydrated; stained in 0.1% Fast Green in ethanol; and finally washed, cleared, and placed under cover slips. Fast Green stains collagen blue-green, while PTAH stains muscle striations purple-blue and many tissues (including collagen) various shades of red-pink. When we double-stained for collagen and muscle, the pink PTAH pigments were washed out with ethanol and the collagen was re-stained with Fast Green to achieve darker and more distinct color. Each slide-mounted section of a whole pad was placed under a Zeiss Opmi 6-CFC dissecting microscope. Photomicrographs were taken at 8x magnification with a Canon Digital Rebel camera.

### Overview of a whisker deflection experiment ex vivo

To constrain some model parameters, we used data from a study that experimentally quantified tissue deformation in an *ex vivo* preparation [29]. In these experiments, the displacement of tissue internal to the follicle near the RS level is imaged during external deflection of the whisker. An overview of the experimental procedure is provided here.

Briefly, Whiteley et al. dissected the C-row of whiskers, suspended C1 horizontally in a petri dish, with flanking follicles supported by silicone, and deflected the C1 whisker proximally (~7mm) and horizontally with a high-resolution manipulator (Fig 1A). The wall of the actuated follicle at the level of the ringwulst was dissected, providing a window (~1×1mm^2^) for imaging the relative displacement of fluorescently labeled Merkel cells. Relative displacement was calculated as the difference between pre-deflection position of a Merkel cell, and its position at the peak of whisker deflection, from z-stacks of two-photon images. Displacement was considered using a cylindrical coordinate system within the follicle consisting of 3 dimensions: radial (displacement perpendicular to the whisker), longitudinal (displacement along the whisker), and polar (rotation around the whisker, clockwise from dorsal axis being 0°) (Fig 1B). Whiteley et al.’s results indicated that the whisker at the RS level moves to the opposite side of the follicle, in the direction of deflection. Total displacement in the three dimensions was 4.8μm. Finally, no sign change of radial displacement was identified in the observed window.

**Fig 1.**
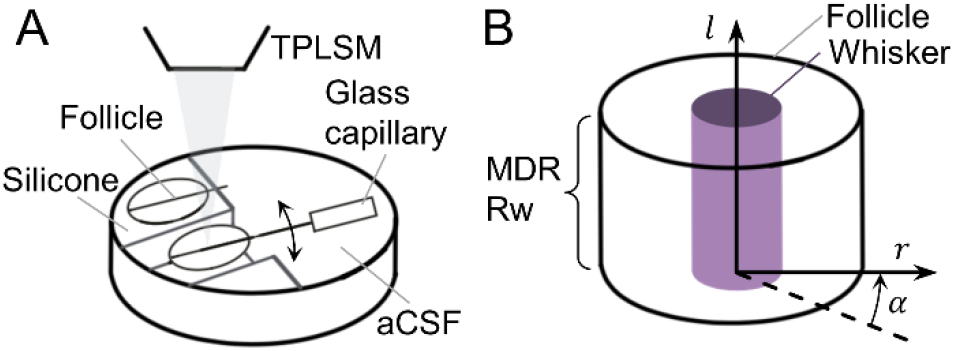
Schematic of the experimental procedure and coordinate systems used in Whiteley et al. [29]. (A) A row of follicles pinned to silicone base, immersed in artificial cerebrospinal fluid for two-photon imaging. A single whisker was deflected by displacing the glass capillary in rostral or caudal directions, while the dorsal surface of the follicle was exposed and imaged. TPLSM: two-photon laser scanning microscope; aCSF: artificial cerebrospinal fluid. (B) A cylindrical coordinate system was used for displacement analysis of the tissue between the whisker (purple cylinder) and the follicle. A section of the Merkel cell dense region at the level of the ringwulst was imaged. Radial distance (r) measures displacements perpendicular to the vibrissa; polar angle (α) measures displacements around the circumference of the vibrissa; and longitudinal distance (l) measures displacements along the vibrissa length. MDR: Merkel-cell dense region; Rw: ringwulst.

### A beam-and-spring model for the vibrissa displacement in the follicle sinus complex

We created a beam-and-spring model (Fig 2A) to simulate deformation of the vibrissa in the follicle and the follicle in the tissue. Two beams and six springs were used.

**Fig 2.**
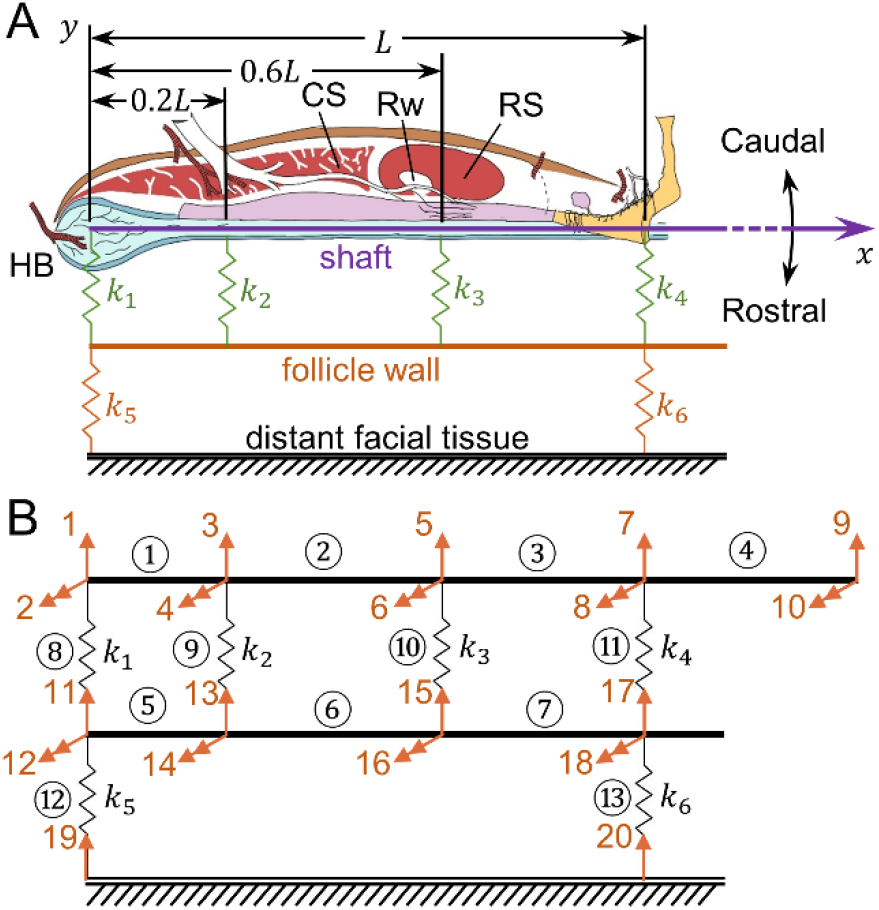
The mechanical formulation of the beam-and-spring model for the FSC. (A) The beam-and-spring model of the vibrissa and tissue. The tissue inside the follicle is modeled by four internal springs (k_1_, k_2_, k_3_, and k_4_), colored in green and placed at anatomically relevant locations. The tissue outside the follicle is modeled by two external springs (k_5_ and k_6_), colored in orange. The vibrissa is represented as a purple line, and the follicle wall is an orange line. The follicle system is supported by the rest of the tissue on the face, considered to be far away and thus indicated as ground. The overall length of the follicle is L, from the base to apex. (B) The finite element definition of the vibrissa-follicle structure. The entire structure is modeled by 13 elements denoted by circled indices, with a total of 20 degrees of freedom (DOFs) denoted by orange indices. Single headed arrows denote displacements; double-headed arrows denote rotations. Elements 1~4 represent the segments of the vibrissa, elements 5~7 represent the segments of the follicle wall, and elements 8~13 represent six springs.

The whisker is represented by a Bernoulli-Euler beam [30] and the follicle wall by a rigid beam. The tissue distribution internal to the follicle wall is modeled by four *internal springs* (k_1_, k_2_, k_3_, and k_4_), chosen at the locations of the hair bulb (HB), the cavernous sinus (CS), the ring sinus (RS), and the follicle entrance. These locations were chosen based on the approximation that material properties are similar within each of the three partitioned regions. The connective tissue and muscle outside of the follicle are represented by two *external springs* (k_5_ and k_6_), representing the locations of the two contact points of the intrinsic muscles at the top and bottom of the follicle, respectively. The adjacent follicles and distant facial tissue are indicated as rigid ground in the schematic. This approximation is appropriate because in all experiments and simulations presented in this work, only a single follicle is deflected at a time. Simulating simultaneous deflection of multiple whiskers is a topic for future work.

All springs are drawn on only one side of the schematic. One should pay attention to the following two points: First, the springs model the net force (either compressive or tensile) in the connective tissue or muscle fiber. Putting springs on one side is mathematically equivalent to modeling the mechanics where elastic tissues are on both sides. Second, the schematic is not intended to imply that the muscle insertion points are only on the edge of the follicle. In fact, the intrinsic muscle generally wraps around the follicle with multiple anchor points.

Although this model is simplified, we emphasize that our goal is only to simulate the overall whisker deformation, not the exact internal tissue displacement and strain. The model captures the essential features of the follicle stiffness distribution and allows examination of a finite, but wide, range of spring stiffnesses. The model is well suited for finite element method analysis. The solution to this problem employs a standard approach of stiffness matrix assembly [31]. A proper decomposition of the structure for finite element analysis is shown in Fig 2B.

With these definitions, we define the global displacement matrix

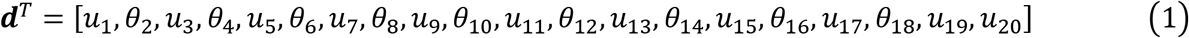

The stiffness matrices for elements 1 to 7 are

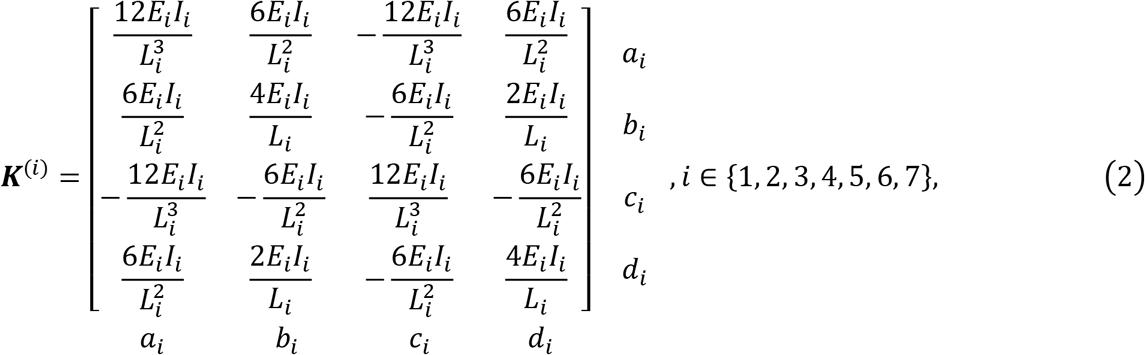

where E is Young’s modulus, I is the moment of inertia (=πR^4^/4, R: vibrissa base radius), and L is the length of the corresponding element. The indices [a_i_, b_i_, c_i_, d_i_] below and to the right of the matrix correspond to the indices of the DOFs for the i-th element when it is added to a global stiffness matrix. For example, [a_1_, b_1_, c_1_, d_1_]=[1,2,3,4] for element 1.

Similarly, the stiffness matrices for elements 8 to 13 are

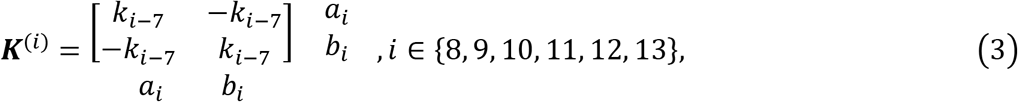

where k_i-7_ are the spring constants for the i-th element. The indices [a_i_, b_i_] below and to the right of the matrix correspond to the indices of the DOFs at each end of the i-th element when it is added to a global stiffness matrix. For example, [a_10_, b_10_]=[5,15] for element 10.

When the global stiffness matrix is constructed, linear shape functions reflect the assumption of constant Young’s modulus along the vibrissa. The global stiffness matrix is a 20-dimensional sparse square matrix, assembled by adding up values from local stiffness matrices sharing the same indices pair defined previously, or by using the scatter operator **L**^(i)^, dependent on indices for corresponding DOFs, to stack the local stiffness matrices to a global sparse matrix. The result is

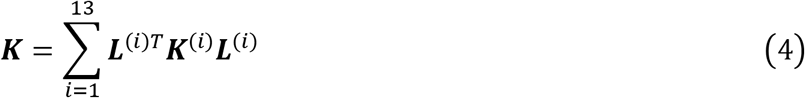

In addition to the natural boundary condition that u_19_=u_20_=0, we also apply the essential boundary condition that θ_8_=−10°, to simulate a 10° deflection of the vibrissa. By using penalty method [32, 33], the overall stiffness matrix and force matrix is given by

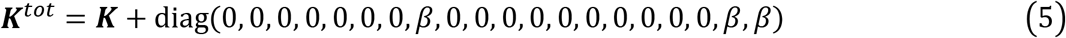

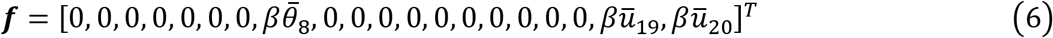

where β is a very large number usually 10^7^ to the average K(ii) to enforce the natural boundary conditions. In our calculation, we choose β=10^7^E (Young’s modulus).

The nodal displacements are then related to the nodal forces by

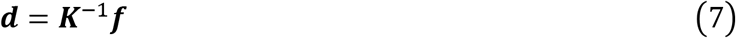

The shear force V(x) along the vibrissa is simply the cumulative sum of the forces caused by the deformation of those springs. For the Euler-Bernoulli beam, the deflection of the beam u(x) is fully described by the equation

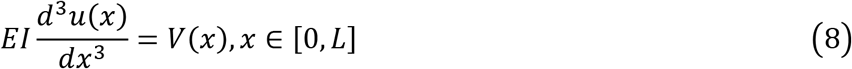

The Young’s modulus (E) for the vibrissa is estimated to be 3.5GPa based on several studies [34, 35], the follicle length (L) is measured to be ~1mm, and the diameter of the vibrissa near its base is assumed to be ~150μm [10]. Note that because the displacements can be expressed in a nondimensional form, change in these estimates simply scales the magnitude of displacements. The boundary conditions are the displacement on both ends and the bending moment on a solved end. In particular, the bending moment for the base of the follicle (free end) is M(0)=0.

Upon deflection of a whisker, the follicle will not stay stationary relative to the animal’s head. We contrast differences and similarities of follicle movement between passive touch and active whisking qualitatively in Fig 3A. The follicle will be driven either by the deflected whisker (passive touch), or by actuated intrinsic muscle (active whisking).

**Fig 3.**
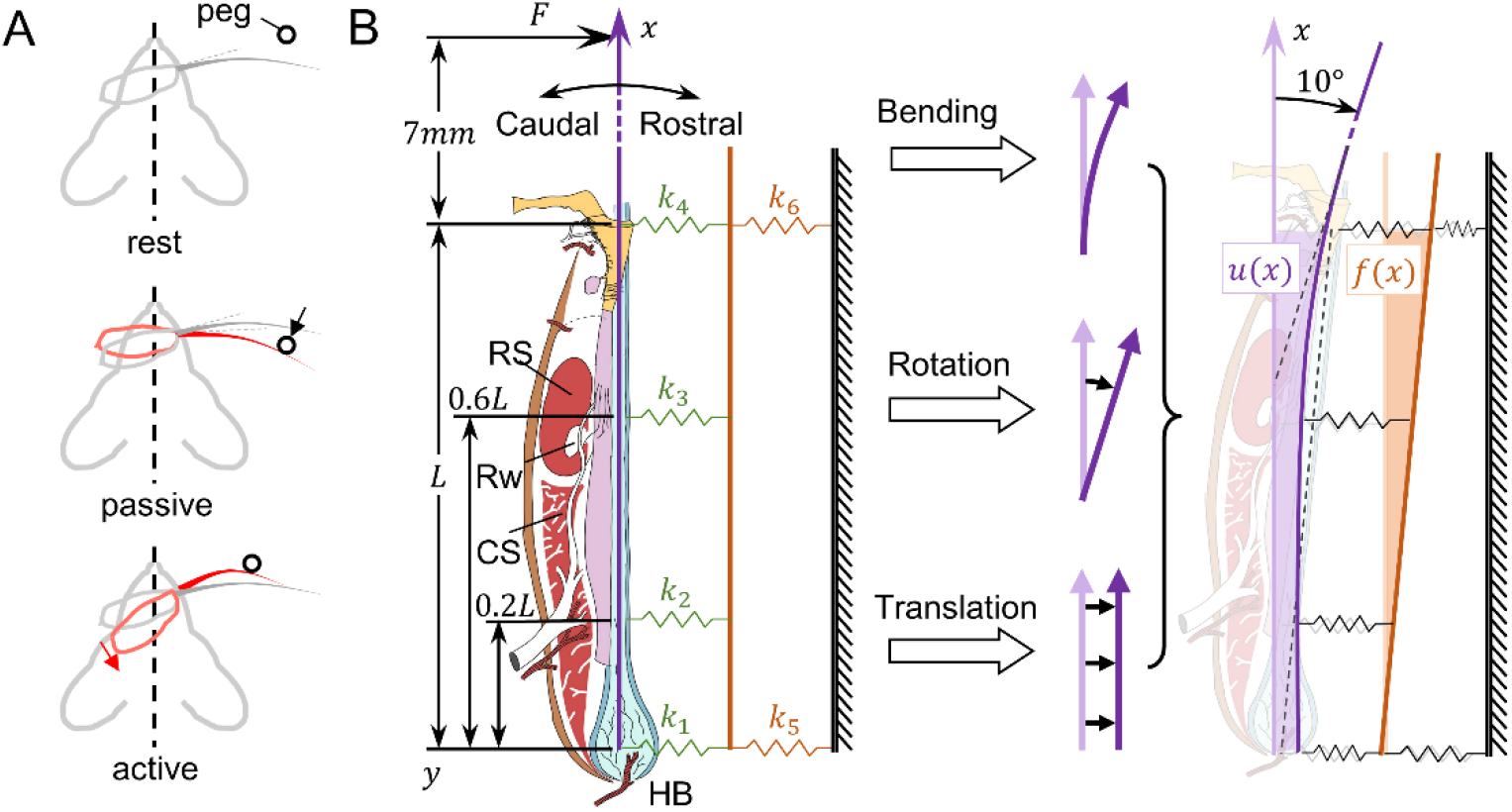
Relative and absolute displacements during whisker deflection. (A) Illustrations demonstrating mechanical differences between passive touch and active whisking. **Top:** the whisker and follicle at their resting locations, undeflected. **Middle**: during passive touch, the whisker is deflected by a peg, leading to movement of the loosely-held follicle. **Bottom**: during active whisking, the follicle is driven by contracted muscle, which is stiff. In both cases, the follicle moves upon deflection. (B) Modeling displacement caused by external deflection of the whisker. **Left**: The beam-and-spring model of the vibrissa and follicle. The undeflected vibrissa lies along the x-axis. Four springs (k_1_, k_2_, k_3_, k_4_) connect the vibrissa (purple) to the follicle wall (orange). Two springs (k_5_, k_6_) connect the follicle wall to distant facial tissue (ground). **Middle**: The deformation of the vibrissa in response to an external applied force F is composed of three elemental components: bending, rotation, and translation. Because the follicle wall is modeled as a rigid beam (i.e., much stiffer than the vibrissa), it displaces only by translation and rotation. **Right**: Schematic after a 10° rostral rotation of the vibrissa. The absolute displacement u(x) is shown in light purple. The follicle wall displacement f(x) is shown in light orange.

We define the following different quantifications of displacement relevant to this study by showing an example of 10° whisker deflection by illustration (Fig 3B). The *absolute displacement* u(x) of the vibrissa is the difference between its deflected position and its original undeflected position for all points on the vibrissa. The deflection of a vibrissa includes bending, rotation, and translation, all of which contribute to the absolute displacement u(x) of the whisker. The *follicle wall displacement* f(x) is similarly defined for the follicle wall. Finally, the *relative displacement* r(x) of the vibrissa is defined as the difference between u(x) and f(x). The relative displacement is more relevant than absolute displacement because it determines how the mechanoreceptors along the whisker length interact with the internal tissue.

## Results

### Parameter constraints and optimization

Although the structure of the mechanical model has been established (Fig 3), the values of the spring constants representing the tissue stiffness are as yet unconstrained. Before discussing potential effects of muscle and blood pressure activity, we impose constraints to bracket the range of spring constants and investigate possible whisker deformation profiles.

#### Skin stiffness imposes constraints on k_4_, k_5_, and k_6_

We begin by constraining k_4_, k_5_, and k_6_. Two external springs (k_5_ and k_6_) represent the muscle attachment and model the rotation and the translation of the follicle within the tissue. When the whisker is deflected passively (e.g., when the animal is anesthetized, resting, or unprepared for an external stimulus), the intrinsic muscle outside the follicle is, by definition, relaxed. This relaxed muscle together with other connective tissue is representative of the overall skin stiffness (8MPa for mouse [36]). Therefore, we approximated the overall skin stiffness in our model as the sum of k_5_ and k_6_ by multiplying the elastic modulus E_skin_ by the follicle length:

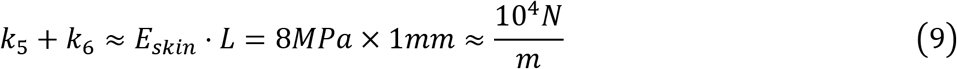

Notice that the ratio of k_5_ and k_6_ will depend on the exact state of the intrinsic muscle; this ratio will become important later in our analysis of active vs. passive deflections.

To constrain the value of k_4_, the spring at the follicle entrance, we noted that previous work has indicated that the vibrissa tends to be stiffly clamped as it enters the follicle [37]. It is clear from the videos associated with this earlier study that the vibrissa displaces very little relative to the follicle at its entrance. To ensure such a rigid vibrissal-follicle junction, k_4_ should be much larger than the unactuated intrinsic muscle stiffness represented by the sum of k_5_ and k_6_. A factor of 100 is sufficient to prevent translation (and permits only rotation) at the follicle entrance:

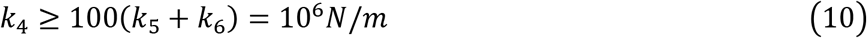

#### Follicle histology imposes constraints on k_1_, k_2_, k_3_, and k_4_

Having established basic constraints for k_4_, k_5_, and k_6_, we next estimated the values for the three remaining internal springs (k_1_, k_2_, and k_3_). To estimate the internal tissue stiffness at different levels of the follicle, we carefully examined serial images of mystacial pad tissue sections featuring follicle cross-sections (Fig 4).

**Fig 4.**
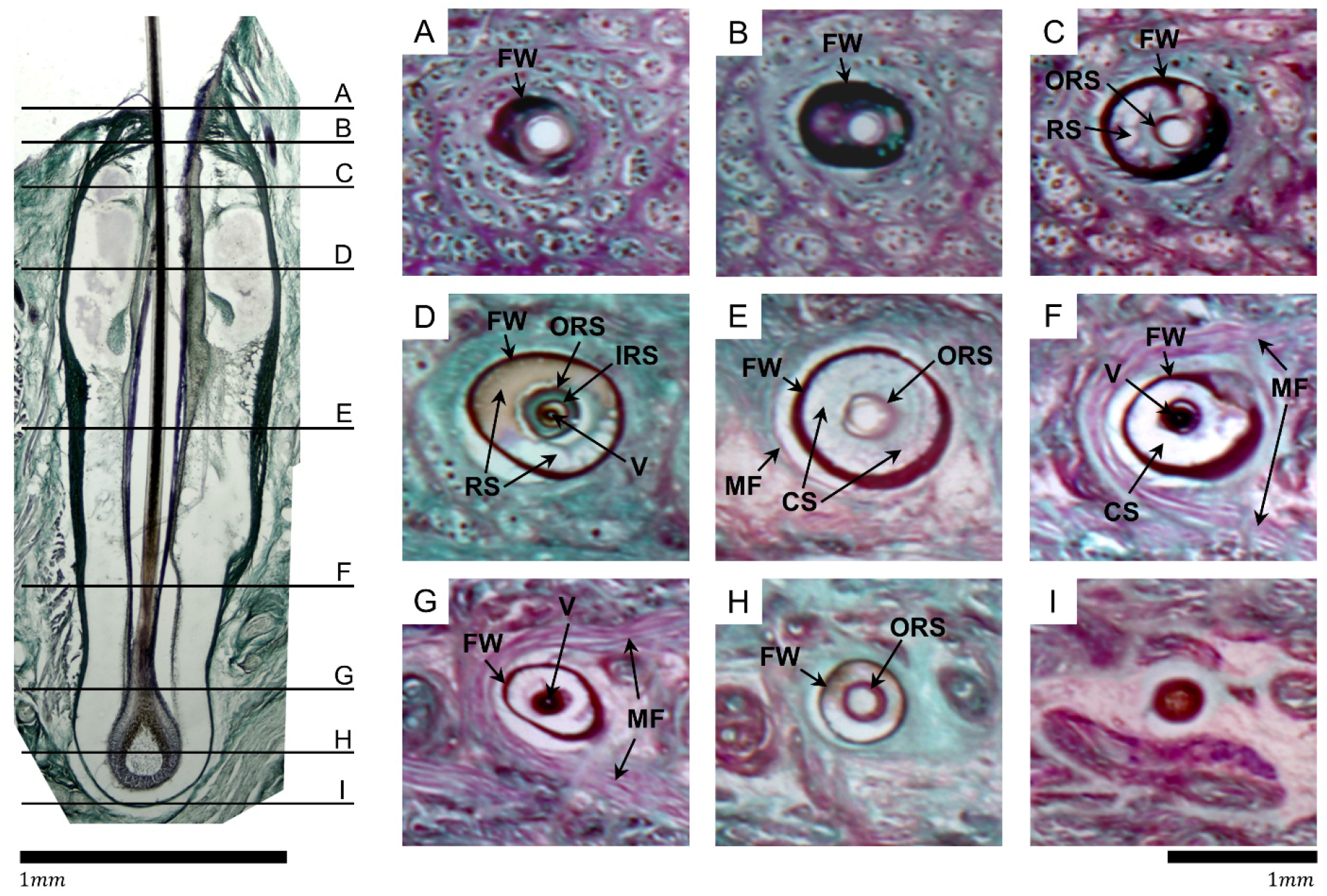
Images of horizontal (lengthwise) and parasagittal (cross-sectional) FSC sections permit estimation of relative tissue stiffness. All images were taken at 4x, and respective scale bars indicate 1mm. The lengthwise image was assembled from multiple tiled images. Brightness, contrast, and magenta (R: 255, G: 0, B: 255) saturation were globally increased during post-processing. The follicle wall is thin at the medial end, thicker in the middle, and thin again towards the apex, before getting much thicker at the insertion into the skin. **Left:** The lengthwise cross-section of the A1 vibrissal FSC, with rostral to the left. The slice does not pass exactly through the central axis, so the vibrissa (the continuous medial dark vertical line) is actually thinner than the full whisker diameter, and no medulla can be observed. Horizontal black lines, labeled A – I, from superficial to deep represent the approximate levels at which the cross-sections (right 3×3 panel) were taken, on an equivalent follicle. **Right:** Cross sections of a single C3 vibrissa from superficial (section A) to deep (section I). The vibrissa is present in sections D, F, G, and H, but has fallen out of the tissue in the remaining sections and is observed as a white oval. The whisker in H is diffuse and appears to be an empty space at 4x, but cells are observable at higher magnification, consistent with other descriptions of the hair bulb and papilla [10]. A, B: a white oval where the vibrissa would be tightly held by the follicle wall (FW) is in the center of the image, and the follicle itself is also held tightly in the skin by the tissue. C: the outer root sheath (ORS), a membrane surrounding the vibrissa, is observed as a dark oval. Because the follicle has been sectioned at a slight angle, the ring sinus (RS, slightly deeper in the follicle) is observable on the rostral side. D: at the level of RS, most of the space inside the follicle is occupied by blood (brown) or empty space (white). The ORS and inner root sheath (IRS) can both be observed as dark ovals. The vibrissa (V) is held closely against the IRS, though the outer layer of the hair shaft is not pigmented so appears very light white/gray. E: Leaving the RS, all internal tissue becomes less stiff, appearing less darkly stained. At the level of cavernous sinus (CS), the FW is very thick compared to the vibrissa. MF: muscle fiber. F: Medially to the trabecula-dense region of the CS the internal membranes become denser (darker) but the vibrissa shaft becomes more diffuse, with melanocytes (pigmented portion of the whisker) no longer segregated to the center of the shaft (more apparent at higher magnification). G, H, I: all surrounding tissue is much less dense towards the end of the follicle, near the HB level. The whisker is also less dense.

Because all tissue sections were processed with identical sectioning and staining techniques, the darkness of stain within a given color channel (red/purple, or green) reflects the density of the major structural proteins (keratin and collagen). Keratin is stained pink, and collagen is stained green. Consequently, the images serve as indicators of relative stiffness at different levels of the follicle. Three major inferences about relative stiffnesses at various levels within the FSC can be made from these images.

First, throughout the length of the follicle, the follicle wall is very darkly stained, indicating that it is stiffer than other comparable tissues (other green structures, also mainly composed of collagen). The follicle wall is relatively thin in section B, increases in thickness through section F, and becomes thinner again starting with section G. Note that the follicle has been sectioned at a slight angle so the follicle wall is a bit thicker on the caudal side; this effect is particularly noticeable in sections C and F. This anatomy validates the modeling assumption made in *Materials and Methods* that the follicle wall is rigid compared to the tissue inside and outside (Fig 3).

Second, the stiffness of the tissue internal to the follicle generally decreases from superficial to deep. At the skin surface (sections A, B, represented by k_4_), the whisker (missing) is seen as a white oval, and is densely surrounded by darkly stained tissue. The absence of the whisker from some sections is an artifact of slide preparation, and occurs because the keratin from which the whisker is composed is highly crosslinked, and so does not have many available binding sites for fixative. Consistent with previous studies [37], we find that the whisker is held tightly at the apex of the follicle. This effect is partially attributable to the intrinsic stiffness of the tissue, but also occurs because a small opening restricts the displacement of the vibrissa at the entrance. These features support the previous assumption that k_4_ should be large.

In section C, the outer root sheath becomes visible as a dark band around the whisker, and the surrounding dense staining indicates that the whisker continues to be held tightly within the follicle. Starting with level D, near the RS (k_3_), the whisker begins to be less tightly held within the follicle, as indicated by the lighter staining between the outer root sheath and the follicle wall. Sections E and F show even weaker staining inside the follicle near the CS (k_2_), indicating a continuing decrease in stiffness. This trend continues through section G, which approaches the plate, a mat of connective tissue that loosely overlies the bone [38]. Finally, in sections H and I (k_1_) we see very diffuse keratinocytes that will become the whisker cortex, and the very end of follicle capsule. In these sections the tissue is more hydrated and less dense, and the whisker is quite loose within the follicle.

Third, the follicle is held fairly tightly within the skin in sections A, B, and C. In section C, the whisker is primarily surrounded by keratin (stained pink), which is less elastic and tougher than collagen. By section D, the surrounding matrix is primarily loosely coiled collagen (stained green) and a large crescent of pale green to white is visible around the follicle’s rostral edge. This lighter staining indicates that the follicle is held loosely in the skin. In section E, the first fibers of the sling muscle are visible as pink strands running across the rostral arc of the follicle, and again a space lies between the follicle wall and the muscle fiber, showing that there is not a lot of connective tissue anchoring the follicle to surrounding tissue. The muscle can slide across the follicle and the follicle can slide around in the skin very easily. Sections G, H, and I, continue this trend, showing large white/light green rings/crescents around the follicle that indicate that it is not well anchored in the skin.

Together, the images of Fig 4 suggest that it is reasonable to assume that k_1_≤k_2_<k_4_, that k_1_≤k_3_<k_4_, and that k_5_<k_6_. We cannot infer anything about the relative values of k_2_ and k_3_.

### The deepest internal spring k_1_ has a negligible effect on deformation near the ring sinus

The previous section constrained values for the two external springs, k_5_ and k_6_, as well as the most superficial spring, k_4_. We have also constrained the relative relations for the internal springs k_1_, k_2_, and k_3_, but have left their exact values uncertain.

To bracket the possible range of whisker deformation profiles we first investigated how k_1_ affects the relative displacement in the RS region. Specifically, we simulated the rostral deflection of a vibrissa, and observed the deformation profile of the whisker for different values of k_1_, under different combinations of k_2_ and k_3_.

Fig 5 illustrates the different possible shapes of the relative displacement given a 10° deflection. The value of k_2_ changes across plots from top to bottom, and four distinct values of k_3_ are indicated as four different colors. For each k_2_ and k_3_ combination, changing k_1_ results in a shaded area bounded by two extreme deformation profiles. The width of the shaded area can be as large as 13.0μm microns near the CS (for the smallest value of k_2_). However, the width of the shaded area at the RS level is very small for all combinations of k_2_ and k_3_ (0.286μm in average), indicating that the effect of k_1_ is negligible in this region. Because our primary area of interest is around the RS in later stages of simulation, we therefore set k_1_ to an intermediate value (10^3^N/m) to reduce the degrees of freedom in subsequent parameter tuning.

**Fig 5.**
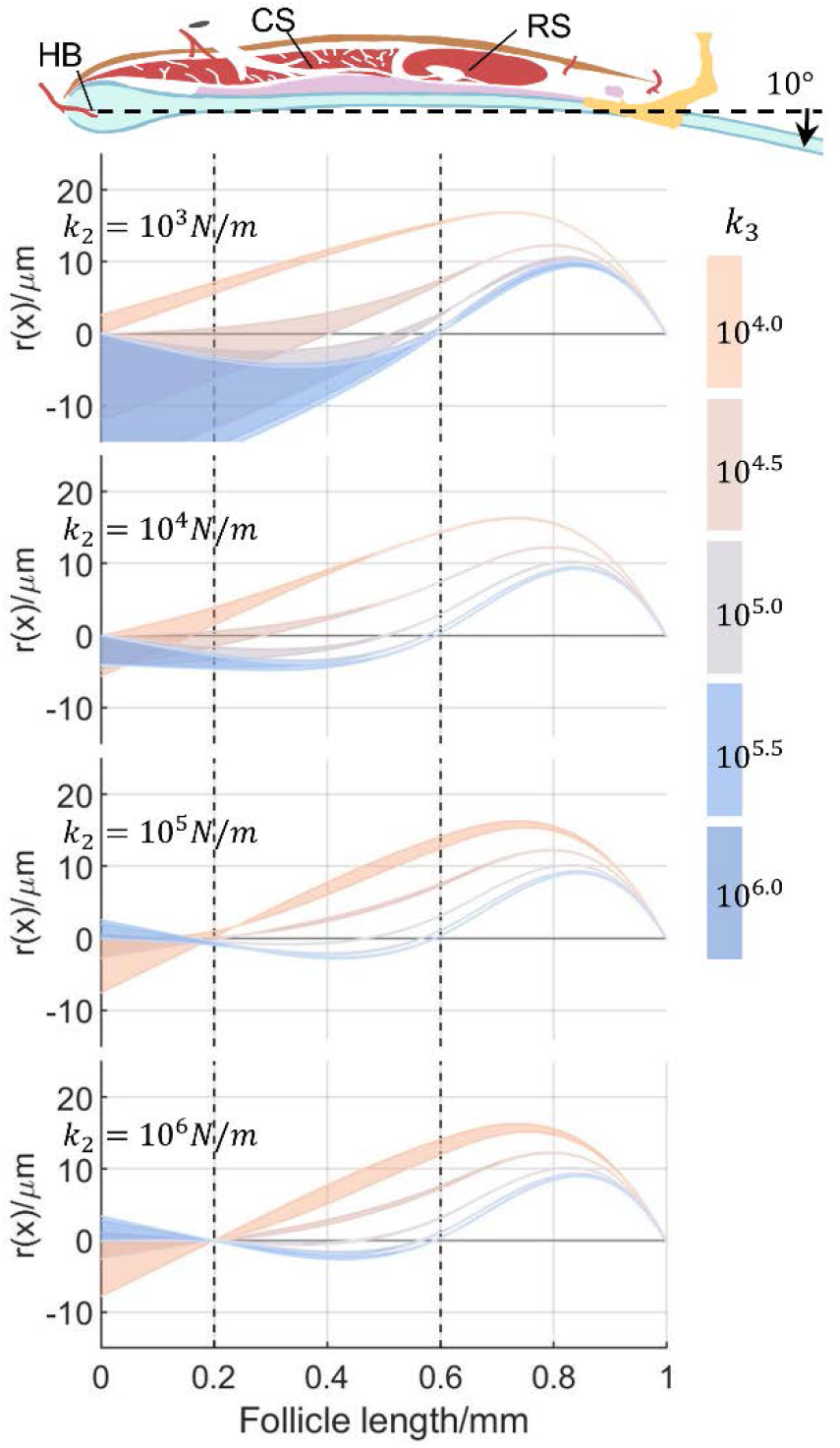
Varying k_1_ has a negligible effect on whisker deformation near the level of the RS. The schematic of the follicle at the top provides a visual reference for anatomical locations. Abbreviations: HB: hair bulb; CS: cavernous sinus; RS: ring sinus. The whisker is simulated to experience a 10° deflection for different combinations of spring constants (k_1_, k_2_, k_3_), and relative displacement r(x) is plotted against the follicle length. The aspect ratio of all plots has been exaggerated. From top to bottom, each of the four plots shows a different value of k_2_, increasing from 10^3^N/m to 10^6^N/m, logarithmically spaced. Within each plot, each color indicates a different value of k_3_, increasing from 10^4^N/m (orange) to 10^6^N/m (blue), logarithmically spaced. Each single shaded area represents the effect of varying k_1_ between 10^2^ and 10^6^N/m, for particular values for the other springs (k_2_ and k_3_). The width of each shaded area can be quite large at the level of the CS especially in the top two panels, but small (0.286μm on average) at the level of the RS.

### The internal spring k_3_ has the largest effect on deformation around the RS

With the value of k_1_ now fixed, we examined the effect of different k_2_ and k_3_ values on the whisker deformation profiles. Depending on different (k_2_, k_3_) pairs, the deformation profile can take different shapes. Fig 6A shows all possible shapes that a vibrissa might take, with three typical shapes indicating three major categories of all deformation profiles, *C-shapes*, *S*_*1*_-*shapes*, and *S*_*2*_-*shapes*. These three different deformation profiles differ by the number of times they cross the x-axis (the resting position). This feature serves as an important indicator to determine the side on which mechanoreceptors are directly exposed to stretched tissue at different levels. For the S_1_-shape (smaller k_2_), the vibrissa shows larger deformation at deeper levels (below the RS). In contrast, for the S_2_-shape (bigger k_2_) the vibrissa is deformed toward different sides of the follicle at different levels, which will in turn affect which groups of mechanoreceptors respond.

**Fig 6.**
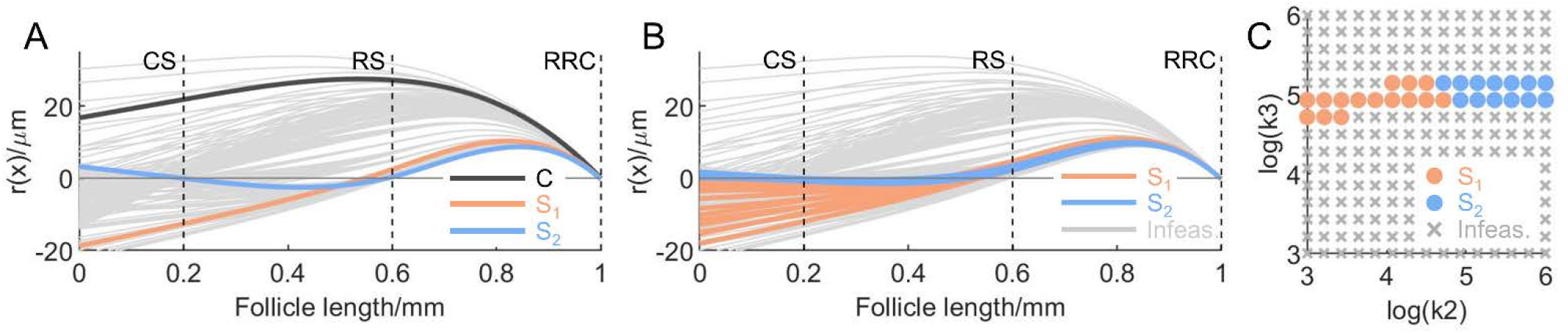
Simulations across all possible (k_2_, k_3_) show that S-shapes are the most probable deformation profiles. (A) Different deformation profiles (r(x)) for different spring pairs (k_2_, k_3_) are shown as a cluster of gray curves. Some typical examples of the deformation profiles of the vibrissa are drawn in black (C-shape), orange (S_1_-shape), and blue (S_2_-shape). (B) Infeasible deformation profiles that violate the experimental data [29] are drawn in gray. Feasible deformation profiles constrained by limiting the displacement at the RS level, r(x)|_x=0.6mm_, to be smaller than 4.8μm are colored in orange and blue, depending on whether it is S_1_- or S_2_-shape. In both (A) and (B) the aspect ratios of all profiles have been exaggerated for visual clarity. (C) Corresponding logarithmically spaced sampling space for (k_2_, k_3_), bounded by [10^3^, 10^3^]N/m and [10^6^, 10^6^]N/m, with similar color scheme. Infeasible space is marked by gray crosses. Feasible space is marked by colored dots.

To further constrain the internal spring constants k_2_ and k_3_, as well as the possible deformation profiles, we noted two features in the experiment by Whiteley, et al. [29]. First, the maximum tissue displacement near the RS relative to the follicle wall is calculated to be 4.8μm (see Methods). For small deflections, the maximum tissue displacement usually takes place at the leading edge (towards the direction of displacement) of the whisker during deformation. We therefore conservatively restricted the whisker displacement at the RS level to be smaller than 4.8μm. Second, there was no sign change of whisker displacement identified in the experiment. We therefore added the constraint that, within the window of the RS (x∈[0.55L, 0.65L]), the whisker must displace entirely towards a single side, opposite to the direction of whisker deflection.

With these two additional constraints imposed, we sampled the spring pair (k_2_, k_3_) drawn from a logarithmically spaced grid bounded by 10^3^N/m and 10^6^N/m in both dimensions, and plotted the relative deformation profiles (Fig 6B). As can be seen, when r(x)|_x=0.6mm_ is constrained to be smaller than 4.8μm, the feasible deformation profile is limited to only the S_1_- or S_2_-shape, excluding the C-shape. Consistent with the second constraint, all feasible profiles in Fig 6B (both S1 and S2) indicate that the tissue is compressed on the side opposite to the deflection direction throughout the region from the RS to the RRC, and stretched elsewhere. This confirms a previous hypothesis [39], that a deflected vibrissa pivots about a fulcrum near the apex of the follicle so that the vibrissa moves in the opposite direction in the RS, causing the tissue in the follicle on the opposite side of the edge to be compressed.

Fig 6C is the corresponding (k_2_, k_3_) sample space, with S_1_- and S_2_-shaped profiles indicated with the same color scheme as in Fig 6B. The feasible space of (k_2_, k_3_) depends only weakly on k_2_, but strongly on k_3_. The value of k_3_ determines the whisker displacement around the RS level, and the value of k_2_ mostly determines whether the whisker takes an S_1_- or S_2_-shape.

### Active touch: Stiffer and more balanced external tissue due to intrinsic muscle activity results in larger whisker deformation during active whisking

We simulated how different muscle activation will affect the relative whisker displacement during both passive deflection and active whisking, for both rostral and caudal deflections. We start by inspecting different possible cases of muscle activities.

Fig 7A shows a simple analysis for the direction of reaction forces exerted by the muscles (not by muscle contraction) on the vibrissa when the vibrissa is deflected. During rostral deflection, both muscles push to support the follicle. Consequently, both muscles are in compression. In contrast, during caudal deflection, both muscles pull the follicle in opposite directions and both muscles are stretched. The direction of the reaction forces is independent of the displacement of adjacent follicles. Note that these reaction forces are not necessarily active muscular opposition to deflection, and likely include a large component of resistance from elastic connective tissue. Regardless, muscles show different mechanical properties depending on their state. Relaxed muscles are like slack “ropes,” yielding almost no stiffness up to a point, and then much larger stiffness when the muscle is stretched to the limit of its constitutive filaments and connective tissue than when at rest or compressed. Contracted muscles are like “springs,” yielding comparable but large stiffness during stretching from the resting length [40].

**Fig 7.**
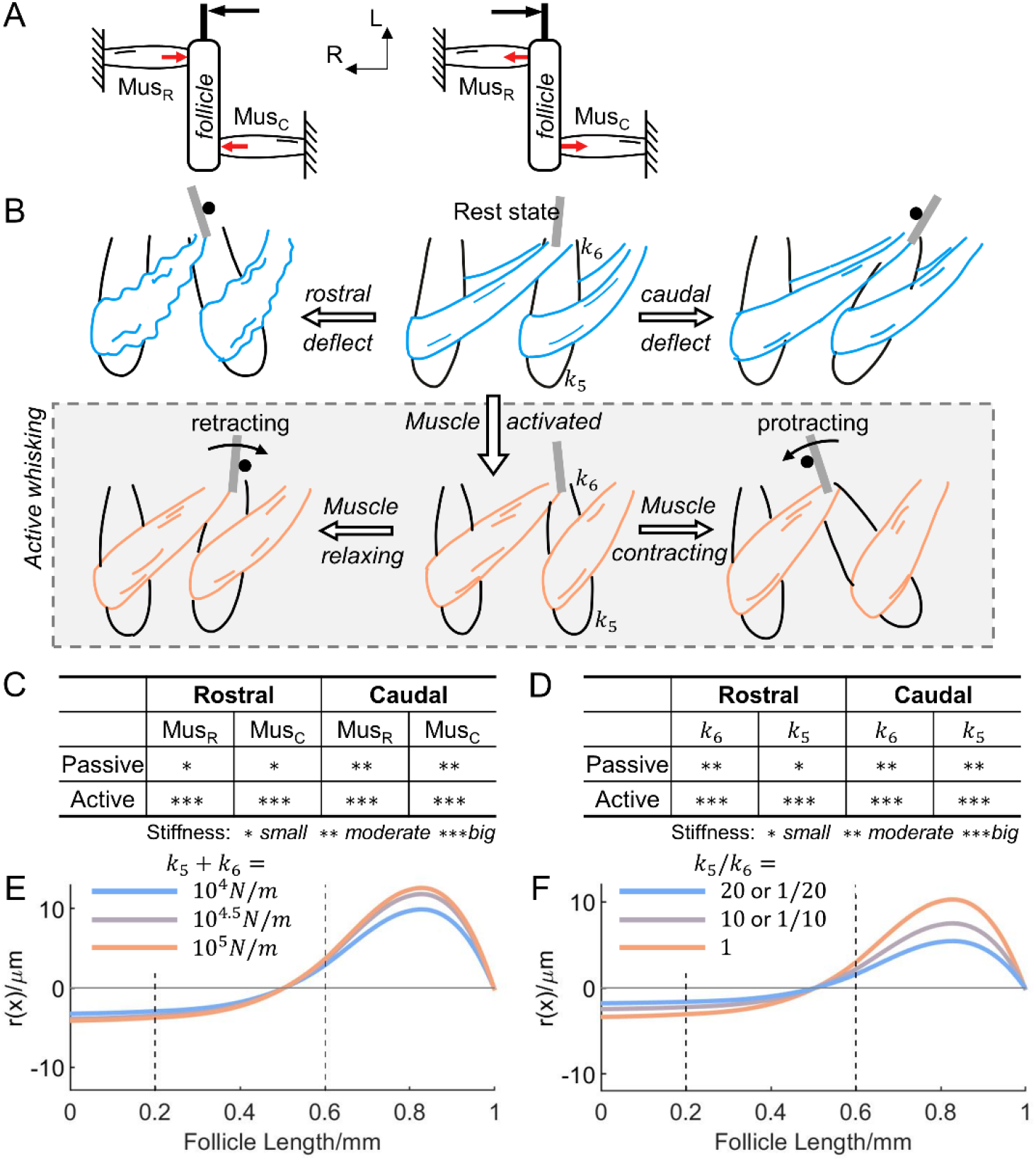
Dorsal deflection and actuated intrinsic muscles result in increased whisker displacement. (A) Illustrations show the directions of the reaction forces on the vibrissa when it is deflected externally. The external force is in black, and the reaction forces are in red. Mus_R_: rostral intrinsic muscle; Mus_C_: caudal intrinsic muscle; R: rostral; L: lateral. (B) Illustrations show the muscle behavior in different circumstances. Only two follicles and the connecting intrinsic muscles are illustrated in one cartoon. One of the two follicles is deflected in either the rostral direction (left column) or caudal direction (right column), during both passive touch (first row) and active whisking (second row, boxed). Muscles are relaxed (blue) for passive touch, but actuated (orange) for active whisking. During passive touch, the whisker is deflected by an external peg, marked by a black circle. During active whisking, the caudal follicle retracts/protracts against a peg, so that the whisker is deflected rostrally/caudally. The locations of two external springs, k_5_ and k_6_, are also indicated. (C) A table summarizes muscle stiffness (in rank order, not proportionally scaled) for different scenarios corresponding to the illustrations. (D) A table summarizes k_5_ and k_6_ stiffness (in rank order, not proportionally scaled) for different scenarios. (E) The change in relative displacement r(x) under different overall external spring stiffness (k_5_+k_6_). The overall stiffness increases logarithmically from 10^4^N/m (blue) to 10^5^N/m (orange). Stiffer external support results in bigger r(x). (F) The identical plot from (E) for different external spring stiffness ratio (k_5_/k_6_). The ratio alters from unblanced (blue) to balanced (orange). More balanced external support results in bigger r(x) at the RS level.

Fig 7B illustrates different scenarios of muscle activity when the vibrissa of one follicle is deflected externally via passive touch (top row) and by active whisking against a peg (bottom row). In all schematics, two neighboring follicles (in the same row) are shown, serially connected by intrinsic muscles [41], and the caudal follicle is assumed to be deflected. During passive touch, the whisker is deflected by a peg in both rostral and caudal directions. During active whisking, the caudal follicle retracts/protracts against a peg in a whisking cycle, so that the whisker is deflected rostrally/caudally. For caudal deflections (rightmost column), both muscles are stretched for passive and active whisking: tensile reaction forces are caused by the peg. For rostral deflections (leftmost column), both muscles are compressed for active and passive whisking: compressive reaction forces are caused by the peg to support the follicle. Based on these schematics and the “ropes” and “springs” analogy, the magnitude of muscle stiffness is summarized qualitatively for different scenarios is summarized in Fig 7C. We emphasize that this summary provides estimates of muscle stiffness, not reaction force. During active whisking there is comparable and large stiffness resisting deflection in both rostral and caudal directions, while relaxed muscles show relatively low stiffness, especially for rostral deflections. It is important to note that during the protraction and retraction phases of an active whisking cycle, the muscles are in a state of either active contraction or passive relaxation, respectively. Though the retraction phase is passive and thus the intrinsic muscles are relaxed, both of these phases have relatively large stiffness due to the existence of extrinsic muscles contributing to greater stiffness in the surrounding tissue.

Next, to assess the possible regulatory mechanism of muscle activity during active whisking, we examined the effects of altering the stiffness of the two external springs, k_5_ and k_6_, in the same scenarios as Fig 7C. These two external springs model the combined stiffness of intrinsic muscles and connective tissue external to the follicle. Similarly, these scenarios of external spring stiffness are summarized in Fig 7D. Specifically, for the top two left entries (rostral deflection during passive touch), the intrinsic muscles are relaxed and compressed, and therefore their low stiffness contributes about as much as that of the connective tissue. As a result of the higher density of collagen and keratin at the skin epidermis, k_6_ is bigger than k_5_. For other entries in Fig 7D, because the connective tissue makes small or negligible contribution to the overall stiffness compared to the intrinsic muscles, the follicle is supported on both ends with about equal stiffness primarily by the muscles. As a result, k_5_ is about the same order of magnitude as k_6_. In the simulation, we examined these effects by changing both the overall stiffness (k_5_+k_6_) and the balance (k_5_/k_6_) of the external support.

We first looked at the overall stiffness of the external support. The overall stiffness is modeled by the summation of the external spring constants (k_5_+k_6_). Fig 7E shows how the relative displacement r(x) changes under different overall stiffness. As expected, stiffer external support in general prevents the follicle from rotating. Consequently, for an imposed external deflection angle, r(x) must increase in magnitude to compensate for the small rotation of the follicle. Specifically, r(x)|_x=0.6mm_, increases from 2.97μm to 3.66μm (123.10% of its original), when the overall stiffness increases from 10^4^ to 10^5^N/m. The result of the simulation suggests that actuated muscles of high stiffness result in larger whisker deformation followed by larger tissue displacement internal to the follicle.

We next looked at the balance of the external support. Fig 7F shows the relative displacement r(x) for different external spring ratios (k_5_/k_6_), with the value of the sum held constant. This result shows that when external support to the follicle is more balanced, the relative displacement of the whisker within the follicle increases. This effect occurs because the follicle itself rotates less when the whisker is deflected (absolute displacement not shown), meaning that the whisker shaft itself must bend more to accommodate the externally imposed movement. Specifically, r(x)|_x=0.6mm_ decreases from 2.97μm to 1.57μm (52.94% of its original), by switching from balanced support (k_5_/k_6_=1) to unbalanced support (k_5_/k_6_=1/20 or 20). The result suggests that more balanced external support would also result in larger whisker deformation, hence larger tissue displacement internal to the follicle.

### Increased blood pressure in the ring sinus results in larger whisker deformation

The blood flow within the follicle at the RS could be regulated by the autonomic nervous system [42]. It has long been postulated that this regulating blood pressure could mediate tactile sensing resolution for certain slowly-adapting receptors [27, 28], allowing animals to have different perceptual sensitivities as needed. However, the relationship between blood pressure in the RS and sensation is difficult to validate *in vivo*. In our model, a change of hydrostatic (blood) pressure in the RS is simulated by changing the value for k_3_, thus allowing us to estimate the effects of changing pressure on whisker deformation within the follicle.

Fig 8A shows that the relative whisker displacement r(x) decreases at the RS level as k_3_ increases, modeling increasing hydrostatic pressure. Specifically, r(x)|_x=0.6mm_ drops from 2.86μm to 0.32μm (11.17% of its original), when k_3_ increases from 10^5^ to10^6^N/m. The general deformation profiles remain similar.

**Fig 8.**
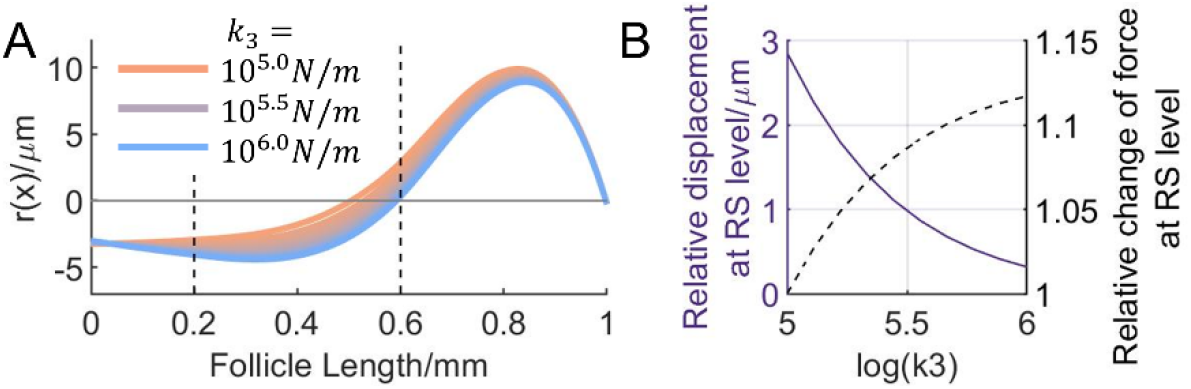
Increasing blood pressure causes decreased whisker displacement, but higher internal forces at the RS level. (A) The change of r(x) as the hydrostatic pressure in the ring sinus is simulated to increase by increasing k_3_. From orange to blue, the value of k_3_ increases logorithmically from 10^5^ to10^6^N/m. With higher blood pressure, r(x) in the RS region decreases. (B) The change in the relative displacement and the force at the RS level as the blood pressure increase. Left y-axis, solid line: relative displacement at the RS level, r(x)|_x=0.6mm_. Right y-axis, dashed line: the relative change of the internal force at the RS level from base state (k_3_=10^5^N/m). The force is given by k_3_∙r(x)|_x=0.6mm_. As the blood pressure increases, the relative displacement at the RS level drops, whereas the force increases.

In the previous section, the whisker displacement is itself a good indicator of internal tissue displacement. Here, due to the inflated and stiffened RS structure, the whisker displacement alone offers only incomplete information about internal tissue deformation. Instead of moving with the whisker in the radial direction, the internal tissue, squeezed between whisker shaft and the RS, could be pushed to other directions. Therefore, we made a preliminary assessment of tissue deformation by looking at the internal force exerted in the tissue at the RS level. This force is defined as the one which occurs between the RS and the whisker shaft, and is simply given by multiplying the tissue stiffness (k_3_) and the relative whisker displacement at the RS level (r(x)|_x=0.6mm_). Fig 8B shows how changes in blood pressure affect relative whisker displacement and the internal force at the RS level, with higher pressure leading to smaller displacement. As blood pressure increases, although whisker displacement decreases, the product of stiffness and displacement increases. In other words, the internal force becomes larger. This force is important if we assume the amount of tissue deformation is related to the force that is applied to it. The implications for mechanosensitivity will be further described in the *Discussion*.

## Discussion

### Advantages and limitations of the beam-and-spring model for the follicle sinus complex

The present work simulated the whisker displacement relative to the follicle during both passive and active deflection, by constructing a mechanical model for the tissue and muscles both inside and outside the follicle. Model parameters were based on an independent *ex vivo* passive whisker deflection experiment and on our own qualitative histological data. The predicted deformation profile is consistent with the hypothesis that the whisker pivots about a fulcrum near its apex, so that the whisker within the follicle (at the level of the RS) is displaced in the opposite direction to the whisker outside of the follicle. Essentially, the tissue in the follicle opposite to the direction of deflection is compressed [39].

By studying the the deformation profile of the whisker within the follicle we have been able to make predictions for which populations of mechanoreceptors will be activated under which circumstances, and to what magnitude. We have calculated whisker displacement profiles along the radial dimension for passive and active whisker contact, including changes in blood pressure in the RS, and intrinsic muscle activity during active whisking. It should be noted that the simulation focuses only on qualitative deformation profiles, and the absolute magnitudes of displacements and forces should not be taken as precise predictions. The limited number of parameters in the current model permits tractable manipulation in order to bracket the range of possible internal mechanical states of the follicle, allowing this first investigation into the effects of internal follicle mechanical parameters on tactile sensory receptor activation and sensitivity regulation.

However, how the mechanoreceptors respond to compressed and/or stretched tissue will depend on the exact details of their shape, location, mechanics, tissue material properties, etc. As the whisker displacement is only an indirect predictor of internal tissue deformation, the model can only predict the relative extent of the mechanoreceptor responses in different regions of the follicle. For example, a larger whisker displacement usually results in larger tissue deformation on the leading edge (towards the direction of displacement), which might further stimulate mechanoreceptors. However, this condition may not apply when other structures inside the follicle (e.g., the blood sinus) are also involuntarily modulated; a small whisker displacement might also accompany a large internal force that stimulates mechanoreceptors when they are pushed laterally.

Additionally, though the current model represents tissue stiffness with springs of different stiffness, they are not directly equivalent. The tissue stiffness is more conventionally represented by material properties such as Young’s modulus. However, it is not appropriate for us to offer precise estimates for Young’s modulus as an accurate prediction of true tissue stiffness at different levels of the follicle. Determining Young’s modulus would require us to assume locations for the borders between areas represented by different springs at a higher spatial resolution than currently possible. The present model is only a first approximation of tissue material properties. In other words, these borders are estimated based on qualitative observation of follicle histology. Fortunately, because the springs are not too densely or sparsely distributed, variations in Young’s modulus should be approximately similar to those of the springs used here.

### Different groups of mechanoreceptors respond to the same direction of deflection

Upon external deflection, the whisker within the follicle displays S-shape deformation patterns (Fig 6). These profiles all share one feature: the part of whisker near the RS displaces in the direction opposite to external whisker deflection, while the part more medial to the RS displaces in the same direction as the external whisker deflection.

Mechanically activated currents have been measured in Piezo2 mechanosensitive channels, one of the mediating channels for Merkel cells’ mechanosensory response [43, 44], during cell-membrane displacements down to 10nm [45]. For the small deflections used in the current simulation and the experiment that inspired it, the electric current in the mechanoreceptors is presumably positively correlated to the cell-membrane displacement. Though little can be inferred from our current model about cell-membrane displacements, we can speculate that mechanoreceptors of a given type found in areas that experience similar tissue deformation are likely to experience similar forces. For example, there are two major groups of Merkel cells inside the follicle, one group densely distributed around the RS level, and one group around the rete ridge collar (RRC) towards the apex. At these two different levels, the whisker displaces to the same direction, causing similar tissue deformation on the same side. Therefore, the model predicts these two groups of Merkel cells will have similar directional responses. These two groups on the same side of the follicle will respond to the same direction of deflection, though which side is not determined. Meanwhile, there may be some populations of receptors that are more activated by stretching or compression. This idea has been confirmed in a recent study [46] in which RS Merkel cells and RRC Merkel cells were found to respond most strongly when the external whisker was deflected in the same direction as the mechanoreceptor. However, it is also important to keep in mind that the present model only applies to quasistatic processes. As Merkel cells are both slowly (RS) and rapidly (RRC) adapting cells [46], these two different groups could also contribute distinct information regarding dynamic processes, which is outside the scope of this study.

### Actuated intrinsic muscles during caudal deflections result in higher tactile sensitivity

Two external springs that represent the external tissue and muscle stiffness are simulated to be actively controlled by the animal during typical active whisking behavior. When the muscle is not actuated, the follicle is weakly supported upon external whisker deflection. More interestingly, during rostral deflection, the follicle also receives a more unbalanced support than during caudal deflection. When a muscle contraction starts, the follicle is strongly supported by actuated muscles, resulting in balanced supports for both rostral and caudal deflection. Our simulation showed that both stiffer and more balanced external support will increase internal whisker displacement (Fig 7). In the case that internal tissue stiffness remains constant, the whisker displacement is positively correlated to the tissue strain field and cell-membrane displacement. This result strongly suggests that caudal external deflection and muscle actuation increase mechanosensitivity; these are precisely the conditions that obtain during active touch.

### Blood pressure in the ring sinus is most likely involved in sensory regulation

Among all internal springs, the one at the RS level, k_3_, is the key parameter in constraining the relative displacement to that observed from the experiment (Fig 6). It is tempting to conclude that this is no coincidence, because the RS is the only structure in the follicle able to adjust its stiffness. Though one role of the RS is to supply blood-born nutrients to cells in the follicle, a change of blood pressure under certain arousal conditions is also postulated to be one of the regulation mechanisms for mechanosensitivity.

Though little can be inferred quantitatively about mechanoreceptor responses at different levels and directions, our simulation showed that a stiffer RS will result in decreased radial whisker displacement, but increased internal force in the radial direction (Fig 8). This internal force is critical to RS-Merkel cells as they are slowly adapting cells [46], making them especially good for sustained pressure sensing. This force might in turn increase the cell-membrane displacement in both polar and longitudinal directions.

### Expected generalizations from passive touch experiments to predictions in active whisking

We noticed that despite regulation by different factors, the deformation profile of the whisker stays qualitatively the same between passive touch and active whisking; the same group of mechanoreceptors will respond when the whisker is deflected in the same direction. This consistency is advantageous to a whisker-specialist animal trying to interpret whisker inputs during both active and passive conditions, because it is easier to interpret a system’s responses if it does not change with behavioral conditions. With that said, the utility of the present model lies in its ability to predict the regions of the follicle in which whisker deformation will activate the mechanoreceptors, even during active whisking. The model offers strong support for possible generalization from passive touch experiments to predictions in active whisking. Future studies that use this model should focus only on the qualitative deformation profiles, but not use any absolute magnitudes of the displacements and forces as precise predictions. The current study paves the way for future studies to more completely characterize internal whisker follicle dynamics to investigate the responses of sub-populations of mechanoreceptors and other specialized compartments within the follicle.

## References

1. Ahl AS. The role of vibrissae in behavior: a status review. Vet Res Commun. 1986;10:245–68.

2. Welker WI. Analysis of sniffing of the albino rat. Behaviour. 1964:223–44.

3. Vincent SB. The function of the vibrissae in the behavior of the white rat. Ph.D. Thesis, The University of Chicago. 1912.

4. Robinson R. A study of sensory control in the rat. Lancaster, PA: The Review Publishing Company; 1909.

5. Kaneko M, Naoki K, Toshio T. Active antenna for contact sensing. IEEE Trans Rob Autom. 1998;14(2):278–91.

6. Solomon JH, Hartmann MJ. Biomechanics: robotic whiskers used to sense features. Nature. 2006;443(7111):525.

7. Birdwell JA, Solomon JH, Thajchayapong M, Taylor MA, Cheely M, Towal RB, et al. Biomechanical models for radial distance determination by the rat vibrissal system. J Neurophysiol. 2007;98(4):2439–55.

8. O’Connor DH, Clack NG, Huber D, Komiyama T, Myers EW, Svoboda K. Vibrissa-based object localization in head-fixed mice. J Neurosci. 2010;30(5):1947–67.

9. Quist BW, Hartmann MJZ. Mechanical signals at the base of a rat vibrissa: the effect of intrinsic vibrissa curvature and implications for tactile exploration. Journal of Neurophysiology. 2012;107(9):2298–312.

10. Ebara S, Kumamoto K, Matsuura T, Mazurkiewicz JE, Rice FL. Similarities and differences in the innervation of mystacial vibrissal follicle-sinus complexes in the rat and cat: a confocal microscopic study. J Comp Neurol. 2002;449(2):103–19.

11. Lottem E, Azouz R. A unifying framework underlying mechanotransduction in the somatosensory system. J Neurosci. 2011;31(23):8520–32.

12. Dehnhardt G, Hyvarinen H, Palviainen A, Gertrud K. Structure and innervation of the vibrissal follicle‐sinus complex in the Australian water rat, Hydromys chrysogaster. J Comp Neurol. 1999;411(4):550–62.

13. Park TJ, Comer C, Carol A, Lu Y, Hong HS, Rice FL. Somatosensory organization and behavior in naked mole-rats: II. Peripheral structures, innervation, and selective lack of neuropeptides associated with thermoregulation and pain. J Comp Neurol. 2003;465(1):104–20.

14. Willingstorfer WJ, Hynek B, Jürgen W. Ovarian growth and folliculogenesis in breeding and nonbreeding females of a social rodent, the Zambian common mole-rat, Cryptomys sp. J Morphol. 1998;237(1):33–41.

15. Hyvarinen H, Kangasperko H, Peura R. Functional structure of the carpal and ventral vibrissae of the squirrel (Sciurus vulgaris). J Zool. 1977;182(4):457–66.

16. Hyvarinen H. On the histology and histochemistry of the snout and vibrissae of the common shrew (Sorex araneus L.). Z Zellforsch Mikrosk Anat. 1972;124(4):445–53.

17. Sarko DK, Rice FL, Reep RL. Elaboration and Innervation of the Vibrissal System in the Rock Hyrax (Procavia capensis). Brain Behav Evol. 2015;85(3):170–88.

18. Marotte LR, Rice FL, Waite PM. The morphology and innervation of facial vibrissae in the tammar wallaby, Macropus eugenii. J Anat. 1992;180:401–17.

19. Reep RL, Stoll ML, Marshall CD, Homer BL, Samuelson DA. Microanatomy of facial vibrissae in the Florida manatee: the basis for specialized sensory function and oripulation. Brain Behav Evol. 2001;58(1):1–14.

20. Sarko DK, Reep RL, Mazurkiewicz JE, Rice FL. Adaptations in the structure and innervation of follicle-sinus complexes to an aquatic environment as seen in the Florida manatee (Trichechus manatus latirostris). J Comp Neurol. 2007;504(3):217–37.

21. Dehnhardt G, Mauck B, Bleckmann H. Seal whiskers detect water movements. Nature. 1998;394(6690):235–6.

22. Hyvarinen H. Diving in darkness: whiskers as sense organs of the ringed seal (Phoca hispida saimensis). J Zool. 1989;218(4):663–78.

23. Stephens RJ, Beebe IJ, Poulter TC. Innervation of the vibrissae of the California sea lion, Zalophus californianus. Anat Rec. 1973;176(4):421–41.

24. Marshall CD, Rozas K, Kot B, Gill VA. Innervation patterns of sea otter (Enhydra lutris) mystacial follicle-sinus complexes. Front Neuroanat. 2014;8:121.

25. Marshall CD, Amin H, Kovacs KM, Lydersen C. Microstructure and innervation of the mystacial vibrissal follicle-sinus complex in bearded seals, Erignathus barbatus (Pinnipedia: Phocidae). Anat Rec. 2006;288(1):13–25.

26. Hyvarinen H, Palviainen A, Strandberg U, Holopainen IJ. Aquatic environment and differentiation of vibrissae: comparison of sinus hair systems of ringed seal, otter and pole cat. Brain Behav Evol. 2009;74(4):268–79.

27. Gottschaldt KM, Iggo A, Young DW. Functional characteristics of mechanoreceptors in sinus hair follicles of the cat. J Physiol. 1973;235(2):287–315.

28. McGovern KA, Marshall CD, Davis RW. Are vibrissae viable sensory structures for prey capture in northern elephant seals, Mirounga angustirostris? Anat Rec. 2015;298(4):750–60.

29. Whiteley SJ, Knutsen PM, Matthews DW, Kleinfeld D. Deflection of a vibrissa leads to a gradient of strain across mechanoreceptors in a mystacial follicle. J Neurophysiol. 2015;114(1):138–45.

30. Hjelmstad KD. Fundamentals of structural mechanics. 2nd ed. New York, NY: Springer Science+Business Media; 2005.

31. Fish J, Belytschko T. A first course in finite elements. West Sussex, England: Wiley; 2007.

32. Saad Y. Interative methods for sparse linear systems. 2nd ed. Philadelphia, PA: Society for Industrial and Applied Mathematics; 2003.

33. Liu JW, George A. Computer solution of large sparse positive definite systems. Englewood Cliffs, NJ: Prentice Hall; 1981.

34. Carl K, Hild W, Mampel J, Schilling C, Uhlig R, Witte H. Characterization of statical properties of rat’s whisker system. IEEE Sens J. 2012;12(2):340–9.

35. Quist BW, Faruqi RA, Hartmann MJZ. Variation in Young’s modulus along the length of a rat vibrissa. J Biomech. 2011;44(16):2775–81.

36. Karimi A, Navidbakhsh M. Measurement of the uniaxial mechanical properties of rat skin using different stress-strain definitions. Skin Res Technol. 2015;21(2):149–57.

37. Bagdasarian K, Szwed M, Knutsen PM, Deutsch D, Derdikman D, Pietr M, et al. Pre-neuronal morphological processing of object location by individual whiskers. Nat Neurosci. 2013;16(5):622–31.

38. Haidarliu S, Simony E, Golomb D, Ahissar E. Muscle architecture in the mystacial pad of the rat. Anat Rec. 2010;293(7):1192–206.

39. Rice FL, Munger BL. A comparative light microscopic analysis of the sensory innervation of the mystacial pad. II. The common fur between the vibrissae. J Comp Neurol. 1986;252(2):186–205.

40. Ralston HJ, Inman VT, Strait LA, Shaffrath MD. Mechanics of human isolated voluntary muscle. Am J Physiol. 1947;151(2):612–20.

41. Dorfl J. The musculature of the mystacial vibrissae of the white mouse. J Anat. 1982;135(1):147–54.

42. Fundin BT, Pfaller K, Rice FL. Different distributions of the sensory and autonomic innervation among the microvasculature of the rat mystacial pad. J Comp Neurol. 1997;389(4):545–68.

43. Coste B, Mathur J, Schmidt M, Earley TJ, Ranade S, Petrus MJ, et al. Piezo1 and Piezo2 are essential components of distinct mechanically activated cation channels. Science. 2010;330(6000):55–60.

44. Lou S, Duan B, Vong L, Lowell BB, Ma Q. Runx1 controls terminal morphology and mechanosensitivity of VGLUT3-expressing C-mechanoreceptors. J Neurosci. 2013;33(3):870–82.

45. Poole K, Herget R, Lapatsina L, Ngo HD, Lewin GR. Tuning Piezo ion channels to detect molecular-scale movements relevant for fine touch. Nat Commun. 2014;5:3520.

46. Furuta T, Bush NE, Yang AE, Ebara S, Miyazaki N, Murata K, et al. The cellular and mechanical basis for response characteristics of identified primary afferents in the rat vibrissal system. Curr Biol. 2020.

